# Transcriptomic profiling reveals a pro-nociceptive role for Angiotensin II in inflammatory bowel disease

**DOI:** 10.1101/2023.06.16.545265

**Authors:** James P Higham, Charity N Bhebhe, Rohit A Gupta, Michael Tranter, Farah M Barakat, Harween Dogra, Natalie Bab, Eva Wozniak, Katie H Barker, Catherine H Wilson, Charles Mein, Tim Raine, James J Cox, John N Wood, Nicholas Croft, Paul Wright, David C Bulmer

## Abstract

**Objective:** Visceral pain is a leading cause of morbidity in inflammatory bowel disease (IBD), contributing significantly to reduced quality of life. Currently available analgesics often lack efficacy or have intolerable side-effects, driving the need for a more complete understanding of the mechanisms causing pain.

**Methods:** Whole transcriptome gene expression analysis was performed by bulk RNA sequencing of colonic biopsies from patients with ulcerative colitis (UC) and Crohn’s disease (CD) reporting abdominal pain and compared with non-inflamed control biopsies. Putative pro-nociceptive mediators were identified based on pathway analysis of differentially expressed genes in IBD tissue and single cell gene expression in colonic neurons. Pro-nociceptive activity of identified mediators was assessed in assays of sensory neuron and colonic afferent activity.

**Results:** RNA sequencing analysis highlighted a 7.6-fold increase in the expression of angiotensinogen transcripts, *Agt,* the precursor to angiotensin II (Ang II), in samples from UC patients (p = 3.2×10^-8^). Consistent with the marked expression of the angiotensin AT_1_ receptor in colonic neurons, Ang II elicited an increase in intracellular Ca^2+^ in capsaicin-sensitive, voltage gated sodium channel subtype Na_V_1.8-positive sensory neurons. Ang II also evoked action potential discharge in high-threshold colonic nociceptors. These effects were inhibited by the AT_1_ receptor antagonist valsartan.

**Conclusion:** Findings from our study identify AT_1_ receptor-mediated colonic nociceptor activation as a novel pathway of visceral nociception in IBD patients with UC. This work highlights the potential utility of angiotensin receptor blockers, such as valsartan, as treatments for pain in IBD.

## Introduction

Despite the marked progress in our understanding of the pathophysiology of inflammatory bowel disease (IBD), including ulcerative colitis (UC) and Crohn’s disease (CD), abdominal pain continues to make a significant contribution to disease morbidity and lowered quality of life. As such, there is an unmet clinical need for the rational development of novel visceral analgesics to treat pain during colitis.

Abdominal pain during colitis develops due to the activation or sensitisation of nociceptors by mediators released from the inflamed bowel. The prolonged activation of gastrointestinal nociceptors leads to the development of visceral hypersensitivity which results in the perception of pain in response to innocuous stimuli, such as bowel movements, and the amplification of noxious stimuli. A more detailed understanding of the mediators responsible for nociceptor activation in colitis and the mechanisms by which they stimulate visceral nociceptors is imperative to facilitating the identification of novel drug targets for the treatment of abdominal pain.

In response to this challenge, we examined gene transcript expression in human colonic biopsies to generate a map of the cell types, signalling pathways and pathophysiological processes underpinning UC and CD, as well as mediators with the potential to stimulate visceral nociceptors (i.e., those for which receptors are expressed by colonic nociceptors (Hockley *et al*., 2019)). This data verified previous reports showing elevated expression of angiotensinogen (Agt) mRNA in the inflamed bowel (Garg *et al*., 2020), particularly in UC (Massimino *et al*., 2021), which we identified as a potential mediator of visceral nociception due to the marked expression of the angiotensin AT_1_ receptor, activated by angiotensin II (Ang II), the major biologically active metabolite of Agt, in putative colonic nociceptors.

Although the circulating Agt and Ang II levels are unchanged in IBD, a marked increase is found within the inflamed bowel (Garg *et al*., 2020), where Ang II concentrations correlate with endoscopically graded bowel inflammation (Jaszewski *et al*., 1990). This local production of Ang II may be driven by Cathepsin G (Dabek *et al*., 2009) which cleaves Agt and Ang I to form Ang II (Owen and Campbell, 1998), though renin and angiotensin converting enzyme are also present in abundance in the intestine. Other leukocytes, such as macrophages, synthesise and release Agt (Gomez *et al*., 1993), providing a source of substrate for Agt-cleaving enzymes in inflamed tissue. Indeed, both stimulation of the renin-angiotensin system and Ang II infusion promote colitis in experimental models (Shi *et al*., 2016), while blockade of the AT_1_ receptor ameliorates colonic inflammation in both mice with experimental colitis (Katada *et al*., 2008; Santiago *et al*., 2008; Shi *et al*., 2016) and humans with IBD (Shi *et al*., 2016; Jacobs *et al*., 2019). Despite an increasingly clear role for Ang II and AT_1_ in inflammation, little is known about their function in pain arising from the viscera. The expression of AT_1_ receptors in the *Mrgprd*^+^ population of non-peptidergic (NP) nociceptors (Usoskin *et al*., 2015; Zeisel *et al*., 2018), and colonic sensory neurons (Hockley *et al*., 2019), points to a role in visceral nociception consistent with recent reports that *Mrgprd*^+^ positive nociceptors are important in nociceptive signalling from the colon in mouse (Bautzova *et al*., 2018). Consequently, the aim of this study was to determine the pro-nociceptive properties of Ang II and AT_1_.

## Methods and materials

For full methods and materials, see Supplemental Information.

### Colonic biopsies and RNA sequencing

Colonic biopsies were taken following legal guardian consent from paediatric patients undergoing colonoscopy as part of their routine medical care. For patient data, see Supplemental Data 1. Biopsies were taken from patients undergoing diagnostic colonoscopy at the Royal London Hospital. Colonic biopsies were taken from sites of inflammation in UC patients, and CD patients divided into drug naïve (CDN) and treatment refractory (CDT) subgroups, with all groups reporting abdominal pain in the 4 weeks prior to endoscopy. Biopsies were also taken from the sigmoid colon of patients who reported abdominal pain in the 4 weeks prior to endoscopy but showed no signs of inflammation on investigation and were subsequently diagnosed as having recurrent abdominal pain (RAP). Finally, biopsies were taken from the sigmoid colon of non-inflamed control patients who reported no abdominal pain in the 4 weeks prior to colonoscopy and showed no evidence of inflammation following endoscopy. Ethical approval for the study was provided by the East London and The City Health Authority Research Ethics Committee (REC# P/01/023). Biopsy samples were collected in modified Krebs/HEPES buffer from which supernatants were taken for study in a separate series of experiments, following which samples were transferred to RNAlater and stored at -80°C until processing for RNA sequencing. RNA sequencing was performed using an Illumina NextSeq500. Transcript abundance was calculated using HTSeq-counts and differential expression was investigated using edgeR. Enrichment analysis was carried out using EnrichR (Kuleshov *et al*., 2016).

### Ca^2+^ imaging

Mouse dorsal root ganglia (DRG) were extracted, enzymatically and mechanically dissociated and plated onto glass-bottomed dishes. Ca^2+^ imaging was carried out using Fluo-4 (Chakrabarti *et al*., 2020). Magnetic sorting of DRG cultures was used to isolate neurons according to previously published protocols (Thakur *et al*., 2014).

### Electrophysiological recording from the lumbar splanchnic nerve

For both multi- and single-unit recording, the colorectum (from splenic flexure to anus) with the associated lumbar splanchnic nerve was isolated and removed. Nerve activity from whole nerve bundles or teased fibres was recorded using glass suction electrodes as described previously (Hockley *et al*., 2016).

## Results

### RNAseq analysis of colonic biopsies provides insight into the mechanisms underpinning IBD

The gene expression profile of colonic biopsies taken from paediatric patients diagnosed with either UC or CD (treatment naïve) were compared to those from non-inflamed controls (Supplemental Data 1) to identify genes elevated in the inflamed bowel (cut off, p ≤ 0.0001, Supplemental Data 2). In UC biopsies, 504 genes were upregulated, while 428 genes were upregulated in CD (Figure 1Ai). 76 upregulated genes were shared between UC and CD. The treatment naïve CD (CDN) cohort was also compared to a treatment refractory CD (CDT) group with inflammation on endoscopy. In colonic biopsies from the CDT group, 329 genes were upregulated compared to non-inflamed controls; only 25 were shared with those upregulated in CDN patients (Supplemental Figure 1A and B; Supplemental Data 2). We also examined genes downregulated compared to non-inflamed biopsies (Figure 1Aii-iii; Supplemental Data 2). 51 genes were downregulated in CDN, compared to 164 in UC, 31 of which were shared between conditions. 106 genes were downregulated in the CDT cohort, and 20 of these were in common with CDN patients. Biopsies taken from patients with recurrent abdominal pain showed no gene upregulation, and only six genes were significantly downregulated (Supplemental Data 2).

**Figure 1:**
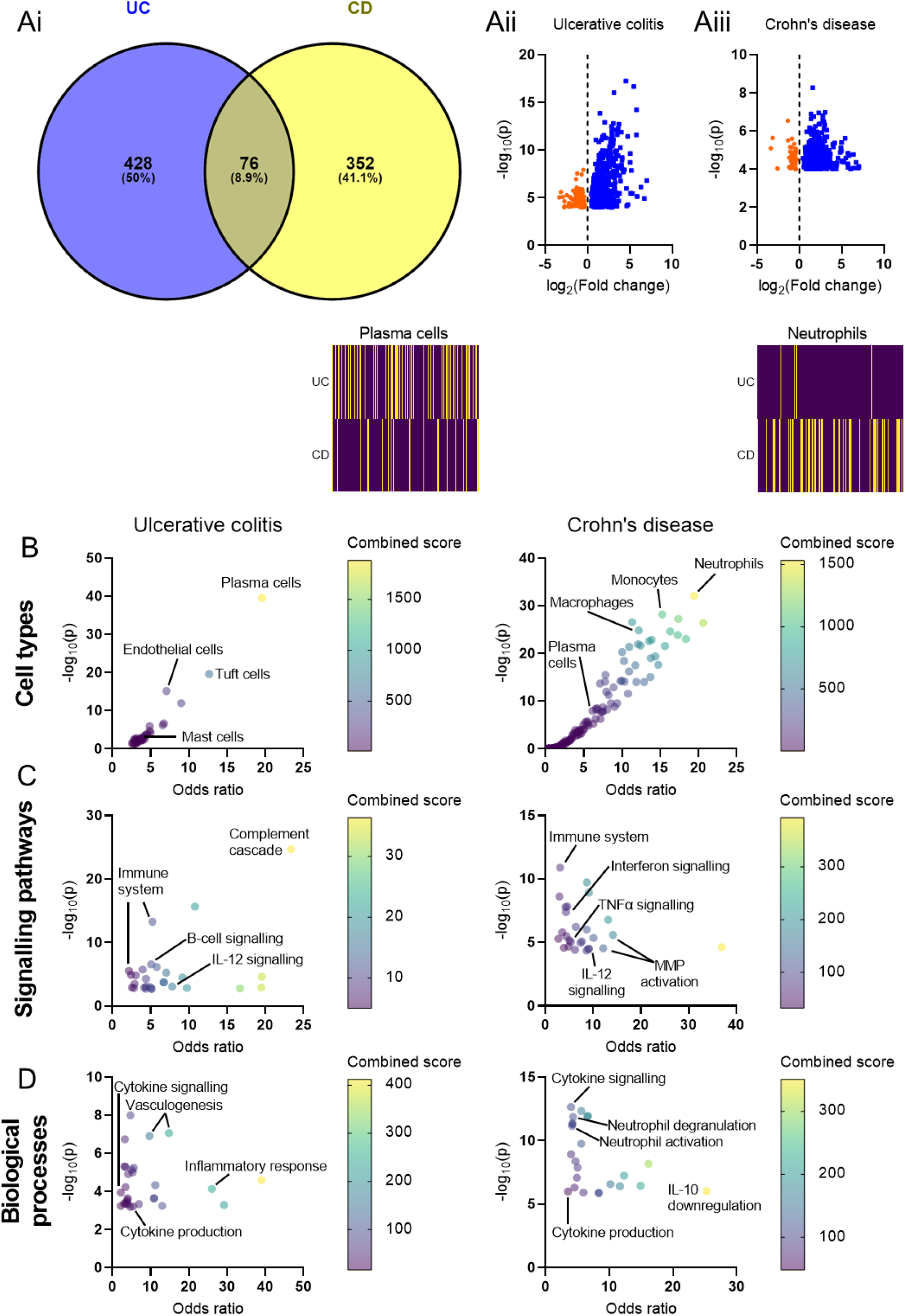
RNAseq and enrichment analysis of colonic biopsies from patients with UC or CD. (A) *(i)* Venn diagram showing the overlap between genes upregulated in UC (blue) and CD (yellow) compared to non-inflamed controls. *(ii & iii)* Volcano plots showing all genes up- (blue) or down- (orange) regulated in UC and CD (p ≤ 0.0001). (B) Bubble plots depicting the enrichment of gene ontology terms corresponding to different cell types in UC (left) or CD (right). Colours represent the combined score, providing a metric of the extent to which genes corresponding to a given ontology term (i.e., cell type) are over-represented (applies to (B) – (D)). *Inset above*: heatmaps showing the annotated gene sets used to identify plasma cells (l*eft*) or neutrophils (*right*); yellow indicates gene upregulation in biopsy tissue. In UC, 52/166 genes annotated to plasma cells were upregulated; in CD, 42/151 genes annotated to neutrophils were upregulated. (C) Bubble plots depicting the enrichment of gene ontology terms corresponding to different signalling pathways in UC (*left*) or CD (*right*). (D) Bubble plots depicting the enrichment of gene ontology terms corresponding to different biological processes in UC (*left*) or CD (*right*).

The upregulated gene sets in UC, CDN and CDT biopsies were compared with annotated gene sets of known biological function (gene ontologies) and enrichment analysis was performed to infer the processes driven by the upregulated genes (Chen *et al*., 2013; Xie *et al*., 2021) (Supplemental Data 3). There was a striking difference in the immune cell types enriched in UC and CDN (Figure 1B). The genes upregulated in UC indicated the elevated presence of plasma cells (p = 2.8×10^-40^), tuft cells (p = 2.5×10^-20^) and mast cells (p = 0.0024); while neutrophils (p = 5.2×10^-35^) and macrophages (p = 5.5×10^-27^) were elevated in CDN (Figure 1B). Given the marked difference in upregulated genes between CDN and CDT biopsies, it is unsurprising that there was also a distinct difference in the cell types present in these groups. There was a reduced enrichment of genes indicating the presence of macrophages and neutrophils in CDT biopsies (Supplemental Figure 1C-E). However, there was an enrichment of various subsets of T cells in CDT biopsies, such as T_memory_ cells (p = 7.6×10^-25^) and T_regulatory_ cells (p = 2.0×10^-19^; Supplemental Figure 1C-E).

The differences in the cell types present are partially reflected in the differential enrichment of signalling pathways (Figure 1C) and biological processes (Figure 1D) in UC and CDN. For example, signatures of B-cell signalling are present in UC (p = 2.8×10^-7^), while signatures of tumour necrosis factor α (TNFα, p = 6.8×10^-6^) and matrix metalloprotease (MMP, p = 2.6×10^-6^) signalling are elevated in CDN. Neutrophils are a major source of both TNFα and MMPs and, consistently, genes associated with neutrophil activation (p = 1.36×10^-12^) and degranulation (p = 4.58×10^-12^) are enriched in CDN (Figure 1D). Signatures of interferon signalling were also identified in CDN biopsies (p = 4.2×10^-8^, Figure 1D).

Enrichment analysis permitted insight into the subcellular compartments involved in signalling events driving UC and CDN. In both diseases, genes associated with secretory granules were enriched (Figure 2A). We therefore sought to identify secreted mediators for which there are receptors expressed on colonic nociceptors. In our biopsy data, *Agt* mRNA was elevated 7.6-fold in UC (RPKM_non-inflamed_ = 1.42±0.17, RPKM_UC_ = 10.83±2.8, p = 3.2×10^-8^) relative to non-inflamed controls (Figure 2B). While the source of *Agt* remains unclear, markers of plasma cells and tuft cells – both highly enriched in UC – correlate with levels of *Agt* across all patient groups (Supplemental Figure 2).

**Figure 2:**
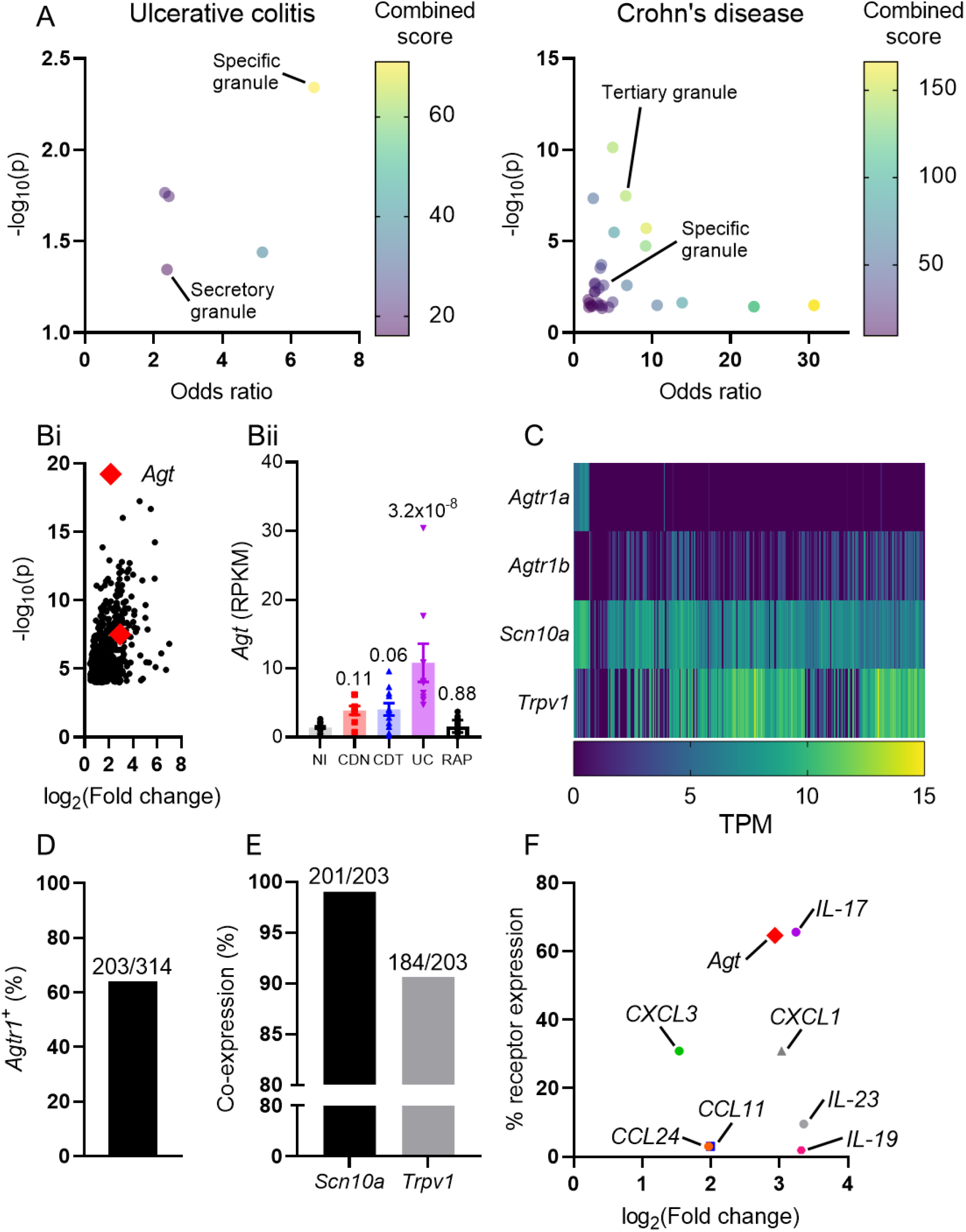
*Agt* mRNA was elevated in UC biopsies, and receptors for Ang II are expressed on colonic nociceptors. (A) Bubble plots depicting the enrichment of gene ontology terms corresponding to different subcellular compartments in UC (*left*) or CD (*right*). (B) *(i)* Scatter plot showing the average fold change (relative to non-inflamed control) for all genes enriched in UC biopsies. *Agt* is highlighted (red diamond). *(ii) Agt* RPKM for each biopsy sample across all patient groups. Benjamini-Hochberg-corrected p-values displayed above bars. (C) Heatmap of gene expression for given genes in mouse colonic sensory neurons. TPM: transcripts per million (expressed as log[TPM]). Data in (C) – (F) redrawn from Hockley *et al*., 2019. (D) The proportion of colonic sensory neurons expressing either *Agtr1a* or *Agtr1b* transcripts. (E) The proportion of *Agtr1*-expressing colonic sensory neurons co-expressing the nociceptive neuronal markers *Scn10a* or *Trpv1*. (F) Scatter plot showing the fold change (relative to non-inflamed controls) for a panel of mediators elevated in UC against the proportion of colonic sensory neurons which express a putative receptor for each mediator.

Receptors for Ang II, the major biologically active metabolite of Agt, are expressed by a large subpopulation of colonic sensory neurons (Figure 2C). 203 of 314 colonic sensory neurons expressed *Agtr1a* and/or *Agtr1b*, the genes encoding the two isoforms of AT_1_ in mouse (Figure 2D, Hockley *et al*., 2019). *Agtr2*, encoding AT_2_, is not expressed by sensory neurons (Usoskin *et al*., 2015; Hockley *et al*., 2019). Of 203 neurons expressing a receptor for Ang II, 201 (99.0%) and 184 (80.8%) co-expressed the nociceptive neuronal markers *Scn10a* and *Trpv1*, respectively (Figure 2E, Hockley *et al*., 2019).

To compare Ang II to other potential targets for investigation, we considered a panel of secreted mediators elevated in UC and examined their fold change and the proportion of colonic sensory neurons expressing a putative receptor for the mediator (Figure 2F). *Agt* was compared to *CXCL1*, *CXCL3*, *CCL11*, *CCL24*, *IL17*, *IL19* and *IL23*. Receptors for CCL11, CCL24, IL-19 and IL-23 are expressed on only a small subset (<10%) of colonic afferents and were not chosen for further investigation. Receptors for IL-17 were expressed by 65.6% of colonic afferents (Figure 2F). It is known that IL-17 interacts with sensory neurons (Richter *et al*., 2012; Segond von Banchet *et al*., 2013), while the effects of Ang II on sensory neurons are less clear and, as such, we have investigated whether Ang II exerts a stimulatory effect on nociceptive sensory neurons.

### Ang II stimulates nociceptive neurons in vitro

To ascertain whether Ang II may be involved in nociceptive signalling, sensory neurons from DRG were cultured and Ca^2+^ imaging was used to determine if Ang II-sensitive neurons were putative nociceptors. A subpopulation of sensory neurons *in vitro* responded to Ang II (2 µM, Figure 3A and B). Ang II evoked a rise in cytosolic Ca^2+^ in 39.8% of sensory neurons (590 neurons from 4 independent cultures), the majority of which were of a small soma area (Figure 3C). Ang II-sensitive neurons had a soma area of 437±16 µm^2^ compared to 821±27 µm^2^ for Ang II-insensitive neurons (p < 0.0001, Figure 3D).

**Figure 3:**
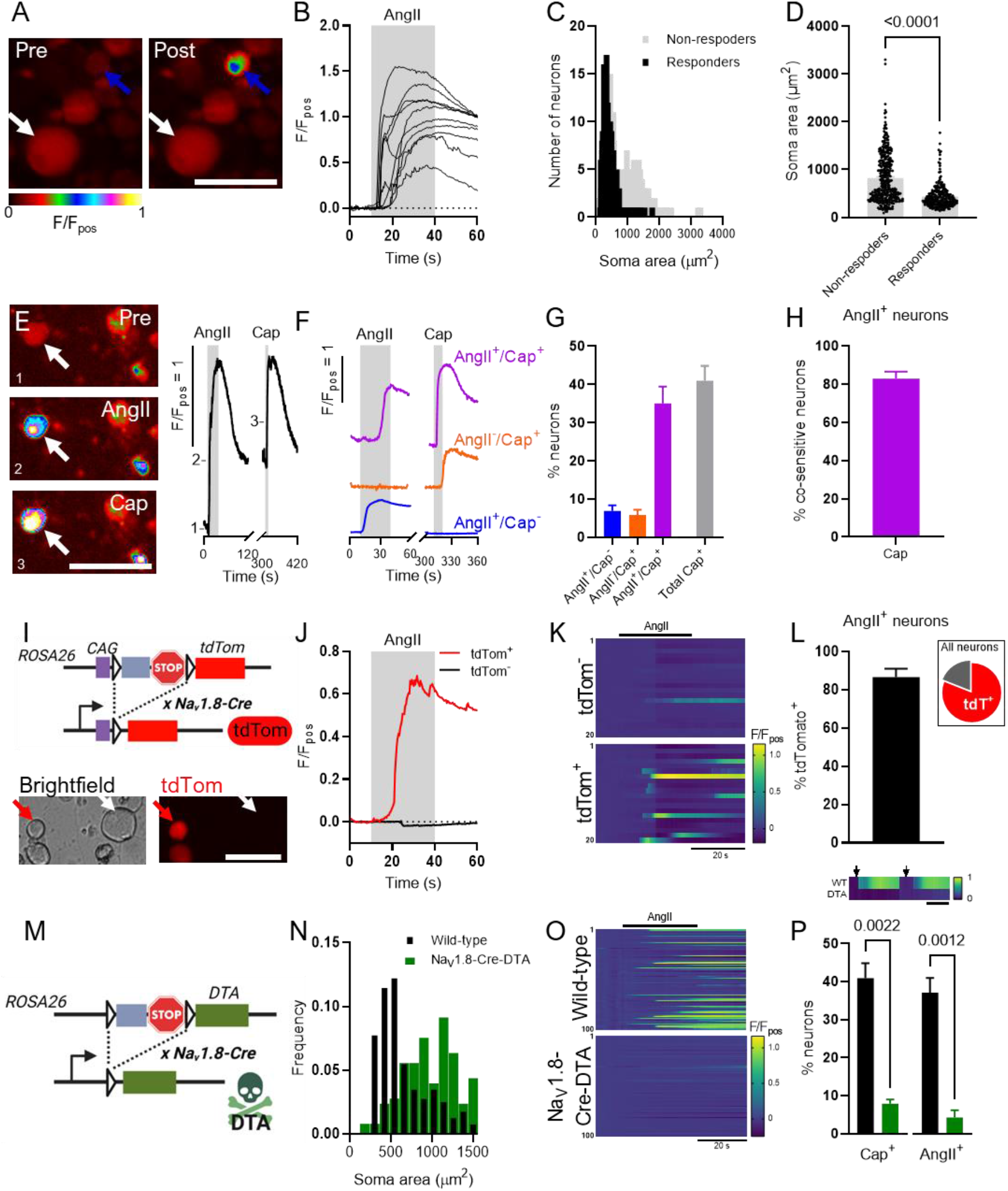
Properties of Ang II-sensitive neurons *in vitro*. (A) False-coloured images depicting Fluo4 fluorescence before (left) and after (right) AngII application. Blue and white arrows highlight exemplar Ang II-sensitive and -insensitive neurons, respectively. Scale bar: 50 µm. (B) Fluo4 fluorescence traces showing the response to Ang II application in 10 randomly-selected AngII-sensitive neurons. (C) Histogram of the neuronal soma area of Ang II-sensitive (black) and -insensitive (grey) neurons. (D) Grouped data showing the neuronal soma area for Ang II-sensitive and -insensitive neurons. 590 neurons (235 Ang II-sensitive and 355 Ang II-insensitive) derived from 4 independent cultures. Data analysed using a Mann-Whitney U-test. (E) (*Left*) False-coloured images depicting Fluo4 fluorescence at baseline (top), during Ang II application (middle) and during capsaicin application (bottom). Scale bar: 50 µm. (*Right*) Fluo4 fluorescence trace showing the response to sequentially applied Ang II (30 s) and capsaicin (10 s) for the neuron highlight by the white arrow (*left*). Numbers on the trace show where the corresponding images on the left were taken. (F) Exemplar Fluo4 fluorescence traces from neurons within each of the populations identified by sensitivity to Ang II and/or capsaicin. (G) Grouped data showing the respective proportional size of each of the identified populations. 221 neurons imaged across 6 coverslips from 4 independent cultures. (H) Co-sensitivity to capsaicin of Ang II-sensitive neurons, derived from the data in (G). (I) (*Top*) Generation of mice expressing tdTomato in Na_V_1.8-positive sensory neurons. A CAG promotor is placed upstream of the floxed (triangles) stop sequence and tdTom to ensure robust expression. A transcriptional stop site halts transcription upstream of tdTom in the absence of Cre. Excision of the floxed stop site by Na_V_1.8-Cre results in the labelling of Na_V_1.8-expressing neurons with tdTomato. Subsequent data (J-L) derived from 384 neurons from 3 independent cultures. (*Bottom*) Images showing exemplar tdTomato^+^ and tdTomato^-^ neurons. Scale bar: 30 µm. (J) Fluo4 fluorescence traces showing the response to Ang II for the neurons highlighted by red (tdTomato^+^) and white (tdTomato^-^) arrows in (I, *bottom*). (K) Heatmaps showing Fluo4 fluorescence during the application of Ang II for 20 randomly-selected tdTomato^-^ (*top*) and tdTomato^+^ (*bottom*) neurons. (L) Proportion of AngII-sensitive neurons which expressed tdTomato and, hence, Na_V_1.8. (*Inset*) The proportion of all neurons imaged which expressed tdTomato (red). 384 neurons imaged across 5 coverslips from 3 independent cultures. (M) Generation of mice expressing DTA in Na_V_1.8-positive sensory neurons. Excision of the floxed stop site with Na_V_1.8-Cre results in the ablation of Na_V_1.8-expressing neurons. Subsequent data (N-P) derived from 312 neurons from 4 independent cultures (wild-type) and 183 neurons from 2 independent cultures (Na_V_1.8-Cre-DTA). (N) Histogram of neuronal soma area for neurons derived from wild-type (black) and Na_V_1.8-Cre-DTA-expressing (green) mice. (O) Heatmaps showing Fluo4 fluorescence during the application of Ang II for 100 randomly-selected wild-type (*top*) and Na_V_1.8-Cre-DTA (*bottom*) neurons. (P) Grouped data showing the proportion of neurons responding to capsaicin or AngII in sensory neuron cultures derived from wild-type (black) or Na_V_1.8-Cre-DTA (green) mice. Wild-type: 312 neurons imaged across 5 coverslips from 4 independent cultures; Na_V_1.8-Cre-DTA: 183 neurons imaged across 6 coverslips from 2 independent cultures. Data analysed using two-tailed Mann-Whitney U-tests. (*Inset, above*) Exemplar heatmaps showing the response of neurons from wild-type (WT) and Na_V_1.8-Cre-DTA (DTA) cultures to the sequential application of AngII and capsaicin (arrows). Scale bar: 30 s.

Sensitivity to capsaicin, an agonist of the non-selective cation channel TRPV1, is a hallmark of a subset of nociceptive neurons. Sequential application of Ang II and capsaicin (1 µM) was used to establish the co-sensitivity between these compounds (Figure 3E). Three responsive populations were identified: those which responded to both Ang II and capsaicin, those which responded to Ang II alone, and those which responded to capsaicin alone (Figure 3F and G). Within the Ang II-sensitive population, 82.8±3.8% of neurons were co-sensitive to capsaicin (Figure 3H) and Ang II-sensitive neurons accounted for 84.9±3.4% of the capsaicin-sensitive population. The response to capsaicin was not sensitised, nor the proportion of capsaicin-sensitive neurons increased, by pre-incubation with Ang II (data not shown), indicating that co-sensitivity was not artificially elevated by the experimental protocol.

The voltage-gated Na^+^ channel, Na_V_1.8, is a key marker of nociceptive sensory neurons. To test whether Ang II-sensitive neurons expressed Na_V_1.8, we first labelled these neurons using Cre-dependent expression of tdTomato (Figure 3I) and investigated the colocalisation of Ang II-evoked Ca^2+^ signals with tdTomato (Figure 3J). Ang II preferentially stimulated Na_V_1.8-positive neurons (Figure 3K): 86.6±4.4% of AngII-sensitive neurons expressed tdTomato (Figure 3L). Of 384 neurons (from 3 independent cultures) imaged, 311 (81.0%) were tdTomato^+^ (soma area: 483±15 µm^2^) and 73 were tdTomato^-^ (soma area: 832±69 µm^2^) (Figure 3L inset), in agreement with previous observations (Thakur *et al*., 2014; Usoskin *et al*., 2015; Luiz *et al*., 2019).

To further ratify the stimulation of Na_V_1.8-expressing neurons by Ang II, these neurons were ablated by Cre-dependent expression of the Diphtheria Toxin A Chain (DTA, Figure 3M). Expression of DTA in Na_V_1.8-positive neurons led to a paucity of small-sized sensory neurons *in vitro* (Figure 3N) and a marked reduction in the response to Ang II (Figure 3O). In wild-type cultures, 37.1±3.9% of neurons responded to Ang II, whereas only 4.2±1.9% responded in cultures from Na_V_1.8-Cre-DTA mice (p = 0.0012, Figure 3P). The response to capsaicin was similarly attenuated (Figure 3P). These data provide functional evidence that Ang II-sensitive sensory neurons *in vitro* exhibit key features of nociceptors.

### AT_1_ is required for the neuronal response to Ang II

There is a clear neuronal response to Ang II in *in vitro* sensory neurons from DRG. Sensory neuron cultures derived from DRG cannot help to resolve whether Ang II directly interacts with neurons because DRG cultures contain myriad non-neuronal cells (Thakur *et al*., 2014). We used magnetic-activated cell sorting (MACS (Thakur *et al*., 2014; Tewari *et al*., 2020)) of DRG cultures to remove non-neuronal cells (Figure 4A). MACS removed non-neuronal cells, identified by positive DAPI staining and negative βIII-tubulin staining (Figure 4B). In unsorted (control) cultures, neurons accounted for 20.2±1.3% of all cells present, compared to 83.8±5.4% after MACS (p < 0.0001, Figure 4C). As reported previously, MACS resulted in a loss of large-sized sensory neurons (see Thakur *et al*., 2014) (Figure 4D).

**Figure 4:**
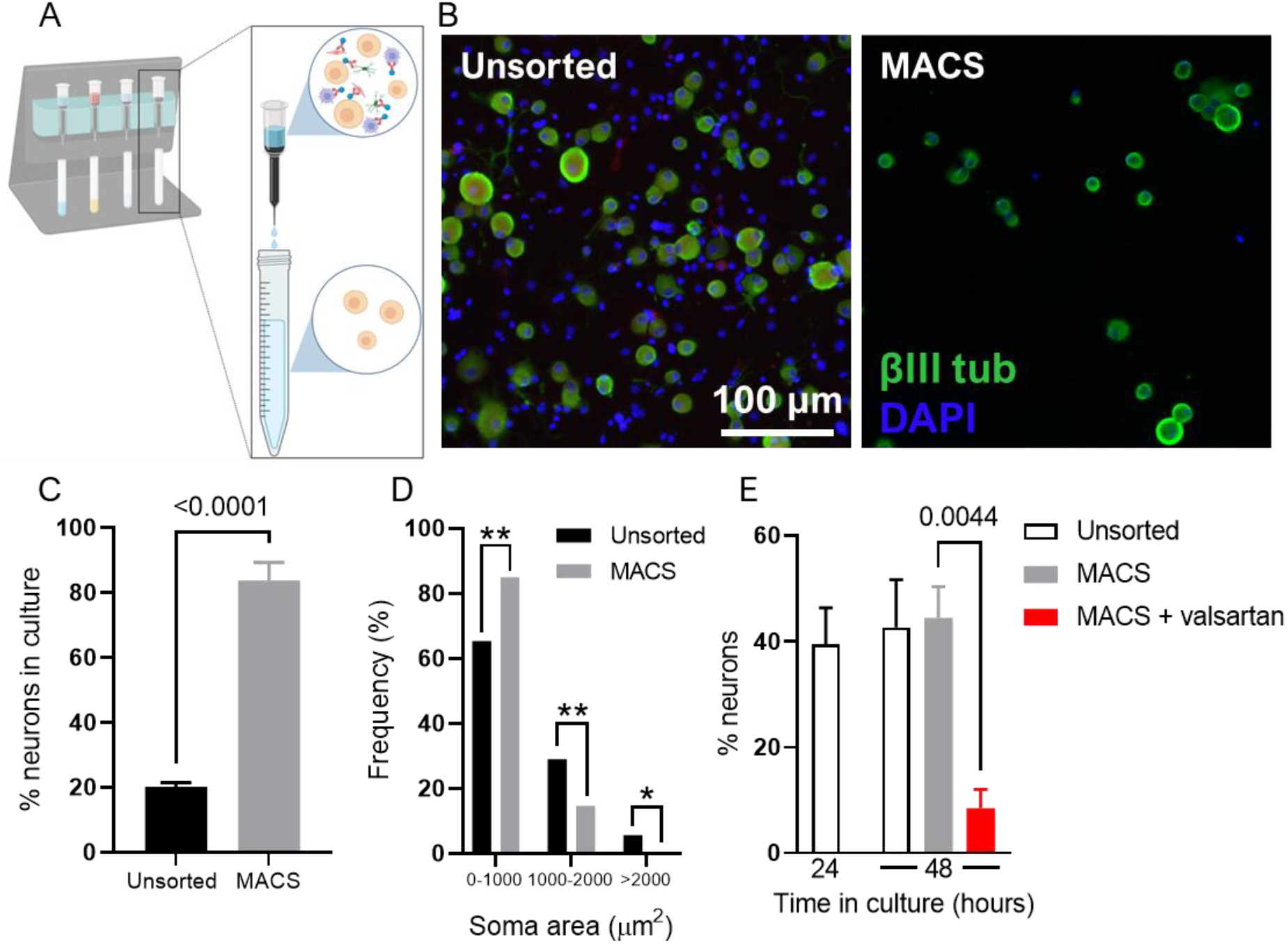
AT_1_R mediated the neuronal response to Ang II *in vitro*. (A) Schematic depicting MACS. Filter columns are loaded onto a magnetic stand (*left*) and the suspension of DRG-derived cells (with non-neuronal cells bound with magnetic beads) is passed through. Non-neuronal cells are retained in the column and neuronal soma are eluted (*right*). (B) Images showing staining for βIII-tubulin (green) and DAPI (blue) in unsorted (*left*) and MACS (*right*) neuronal cultures. A loss of non-neuronal (DAPI^+^/βIII-tubulin^-^) cells is observed. (C) Grouped data showing the proportion of neurons in unsorted and MACS cultures. Unsorted: 9 coverslips from 3 independent cultures; MACS: 11 coverslips from 4 independent cultures. Data analysed using two-tailed Mann-Whitney U-test. (D) Absolute frequency of neurons of given soma areas in unsorted and MACS cultures. Sorting of neurons led to an enrichment of smaller-sized neurons at the expense of neurons >1000 µm^2^. Unsorted: 298 neurons; MACS: 239 neurons. Data analysed using Chi-square tests. (E) Grouped data showing the proportion of neurons responsive to AngII application. Unsorted (24 hours): 5 coverslips from 4 independent cultures; unsorted (48 hours): 6 coverslips from 6 independent cultures; MACS: 10 coverslips from 9 independent cultures; MACS + valsartan: 5 coverslips from 5 independent cultures. Data analysed using a one-way ANOVA with Bonferroni post-tests.

After MACS, sensory neurons required 48 (rather than 24) hours, in culture to properly adhere. This did not have any effect on the response to Ang II (p = 0.99, Figure 4E). In unsorted cultures, 42.7±6.9% of neurons exhibited a rise in cytosolic Ca^2+^ following Ang II application (Figure 4E). After MACS, a similar proportion of neurons responded to Ang II (44.5±5.9%, p = 0.99, Figure 4E). Pre-incubation of sensory neurons with valsartan (1 µM), an AT_1_ antagonist, abrogated the response to Ang II, with only 8.5±3.5% of neurons responding under these conditions (p = 0.0044, Figure 4E). In unsorted cultures, the AT_2_ antagonist, PD123319 (1 µM), had no effect on the neuronal response to Ang II (p = 0.27), while EMD66684 (100 nM; structurally distinct AT_1_ antagonist) did attenuate the proportion of neurons responding to AngII (p = 0.008). These experiments suggest that at least a component of the neuronal response to Ang II is mediated by a direct interaction dependent on AT_1_ and not AT_2_.

### AngII stimulates colonic afferent activity via AT_1_

We used whole-nerve suction electrode recording of the lumbar splanchnic nerve (LSN) innervating the distal colon to ascertain whether Ang II exerted a stimulatory effect on colonic afferents. Bath application of Ang II induced a concentration-dependent increase in afferent activity (Figure 5A and B). 1 µM Ang II (N = 6) evoked a peak increase in afferent activity of 9.5 ± 2.8 spikes s^-1^ (Figure 5C). The area under the afferent response curve (AUC) provides the total spike discharge after AngII application. Under control conditions, the AUC was 5282 ± 843 spikes (Figure 5D). To test the requirement of AT_1_ for the stimulatory effect of AngII, tissue was pre-incubated with one of two structurally distinct AT_1_ antagonists, either valsartan (10 µM, N = 6) or EMD66684 (10 µM, N = 6). Valsartan reduced the peak increase in afferent activity to 0.48 ± 0.52 spikes s^-1^ (p = 0.0043, Figure 5C) and the total spike discharge to 1041 ± 626 spikes (p = 0.0006, Figure 5D). Similarly, EMD66684 attenuated peak and total spike discharge to 1.9 ± 0.92 spikes s^-1^ (p = 0.014, Figure 5C) and 1399 ± 365 spikes (p = 0.0016, Figure 5D), respectively. Losartan, an inverse agonist of AT_1_, also attenuated the colonic afferent response to Ang II (p = 0.0032, N = 6). PD123319 (10 µM, N = 6), an AT_2_-selective antagonist, did not affect peak afferent firing (p = 0.97, Figure 5C) or total spike discharge (p = 0.13, Figure 5D) evoked by Ang II application. Consistently, an AT_2_-selective agonist, CGP42112 (1 µM, N = 6), failed to elicit any change in LSN activity (Figure 5E). In female mice, both baseline and Ang II-evoked LSN activity were lower compared to males, though the proportional change in activity evoked by Ang II was similar between sexes (Figure 5F).

**Figure 5:**
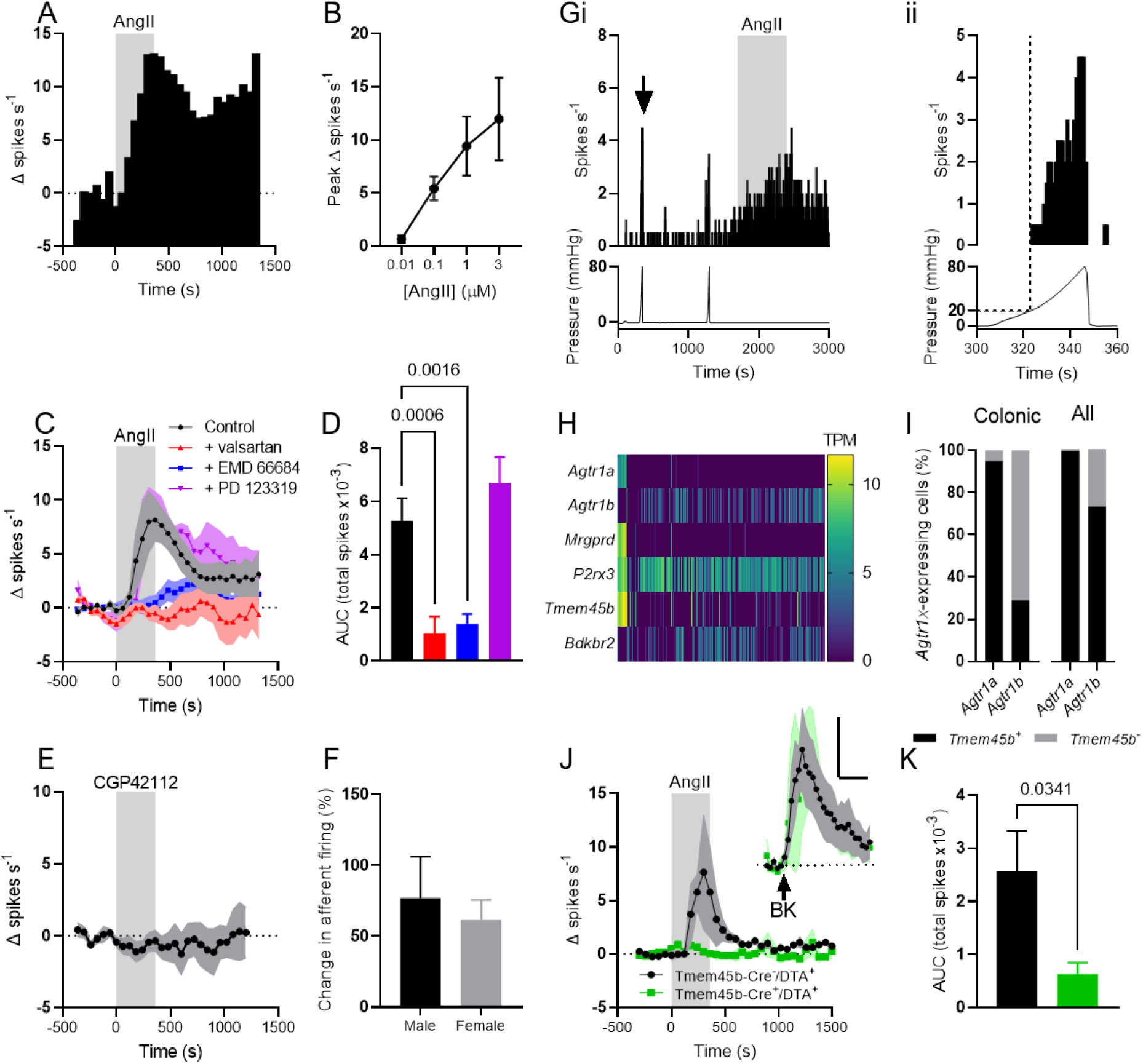
Ang II stimulated activity in the lumbar splanchnic nerve. (A) Exemplar rate histogram showing the change in afferent firing during the bath application of 1 µM AngII (grey shaded area, 0-360 s). (B) Concentration-response relationship for the effect of Ang II on colonic afferent firing. 0.01 µM, N = 4; 0.1, 1 and 3 µM, N = 6. (C) Bath application of Ang II (grey shaded area, 0-360 s) evoked activity in the *ex vivo* LSN under control conditions (black trace). Pre-incubation of tissue with AT_1_ antagonist valsartan (red trace) or EMD66684 (blue trace) blunted the afferent response to Ang II application. PD123319, an AT_2_ antagonist, did not affect the afferent response to Ang II application (purple trace). Symbols show mean change in firing rate (Δ spikes s^-1^) and shaded areas show standard error. (D) Grouped data showing the AUC for the traces shown in (A). Total spike discharge induced by Ang II was attenuated by AT_1_, but not AT_2_, antagonists. Data from 6 independent recordings per condition. Data analysed using a one-way ANOVA with Dunnett’s post-tests. (E) Bath application of CGP42112 (grey shaded area, 0-360 s) did not evoke afferent firing. (F) Grouped data showing the peak percentage change in afferent firing (relative to baseline) evoked by Ang II in tissue from male and female mice. Data analysed using a two-tailed unpaired t-test. (G) *(i)* Exemplar single unit recording showing the response to ramp distension to 80 mmHg and AngII (grey shaded area). *(Top)* Rate histogram of single unit; *(Bottom)* Luminal pressure. *(ii)* Expanded recording showing response to luminal distention marked by arrow in *(i)*. *(Top)* Rate histogram of single unit; *(Bottom)* Luminal pressure. Dotted lines show afferent firing at luminal pressures >20 mmHg. (H) Heatmap of gene expression (log[TPM]) for given genes in mouse colonic sensory neurons, showing co-expression of *Agtr1a* and *Agtr1b* with markers of non-peptidergic sensory neurons, *Mrgprd* and *Tmem45b*. Data redrawn from Hockley *et al*., 2019. (I) *(Left)* Co-expression of *Agtr1a* or *Agtr1b* with the non-peptidergic sensory neuronal marker *Tmem45b* in colonic afferent neurons. Data redrawn from Hockley *et al*., 2019. *(Right)* Co-expression of *Agtr1a* or *Agtr1b* with the non-peptidergic sensory neuronal marker *Tmem45b* in all sensory neurons in the DRG. Data redrawn from Usoskin *et al*., 2015. Black: *Tmem45b*^+^; grey: *Tmem45b*^-^. (J) Bath application of Ang II (grey shaded area, 0-360 s) evoked activity in the *ex vivo* LSN from Tmem45b-Cre^-^/DTA^+^ mice (black trace), but the stimulatory effect of Ang II was reduced in the LSN from Tmem45b-Cre^+^/DTA^+^ mice (green trace). *Inset*: bath application of bradykinin (BK, 1 µM, arrow) evoked similar afferent activity in LSN from Tmem45b-Cre^-^/DTA^+^ and Tmem45b-Cre^+^/DTA^+^ mice. Scale: 500 s, 5 spikes s^-1^. Tmem45b-Cre^-^: 3 mice, 2 male, 1 female; Tmem45b-Cre^+^: 3 mice, 2 male, 1 female. Symbols show mean change in firing rate (Δ spikes s^-1^) and shaded areas show standard error. (K) Grouped data showing the AUC for the traces shown in (J). Data analysed with a one-tailed t-test.

Isolating a small number of fibres from the LSN enabled the analysis of the properties of individual afferent fibres by waveform matching (Figure 5G). 32 individual afferent fibres (from 5 animals) were identified. 16 of the isolated afferent fibres responded to Ang II application, of which 11 were also responsive to colonic distension at pressures >20 mmHg (Figure 5G). In total, 20 of 32 isolated fibres responded to distension pressures >20 mmHg.

Given the co-expression of *Agtr1a/b* with markers of NP colonic afferents, such as *Mrgprd*, *P2rx3* and *Tmem45b* (Figure 5H), we hypothesised that, in mice in which NP afferents had been ablated, the response to Ang II application would be attenuated. We used mice in which Tmem45b-Cre (Figure 5I shows co-expression of *Agtr1a/b* with *Tmem45b*) was used to drive DTA expression, thereby ablating NP sensory neurons (Usoskin *et al*., 2015; Zhao *et al*., 2021). Ang II application resulted in elevated spike discharge in the LSN from Tmem45b-Cre^-^ (control) animals (2575 ± 754 spikes, N = 3, Figure 5J and K). However, the response to AngII was markedly attenuated in Tmem45b-Cre^+^ animals (Figure 5J) with a total spike discharge of 632 ± 213 spikes (p = 0.038, N = 3, Figure 5K). The expression of the B_2_ bradykinin receptor lies mostly outside the NP colonic afferent population (Figure 5H, Hockley *et al*., 2019). Consistently, the afferent response to bradykinin application was not reduced by the ablation of NP neurons (p = 0.48, Figure 5J *inset*).

## Discussion

RNAseq analysis of colonic biopsies provided an insight into the cellular and molecular underpinnings of IBD, ratifying previous observations showing the differential enrichment of immune cell types between UC and CD (Actis *et al*., 2011; Kobayashi *et al*., 2020; Roda *et al*., 2020). UC biopsies were enriched with plasma cells and mast cells, both major effectors of the T_H_2-mediated immune response (Hültner *et al*., 2000; Masuda *et al*., 2002; Maddur and Bayry, 2015). While CDN biopsies were enriched with macrophages indicative of a dominant T_H_1-mediated immune response, in line with signatures of interferon and TNFα signalling. Many genes were upregulated in CDT biopsies, indicative of the presence of multiple T cell subpopulations in keeping with the treatment refractory nature of disease in this patient group. Furthermore, CDT and CDN biopsies differed substantially in their gene expression, suggesting that the bowel in treatment refractory CD patients represents a transcriptional state distinct from both treatment naïve CD patients and non-inflamed patients.

Comparison of differential gene expression for secreted mediators with cognate receptor expression in colonic neurons lead to the identification of Ang II/AT_1_-mediated activation of Na_V_1.8-positive colonic nociceptors as a putative pathway for visceral nociception in UC following elevated *Agt* expression.

This is consistent with work showing correlation of Ang II with endoscopically-graded inflammation in IBD (Jaszewski *et al*., 1990) and more recent reports of enhanced expression of *Agt* in colonic biopsies from patients with IBD (Garg *et al*., 2020) (Massimino *et al*., 2021). Furthermore, retrospective studies of IBD patients prescribed ACE inhibitors or angiotensin receptor (AT_1_) blockers (ARBs) revealed they experienced milder inflammation, reduced requirement for corticosteroid treatment and a diminished risk of hospitalisation and surgical intervention (Jacobs *et al*., 2019; Fairbrass *et al*., 2021; Mantaka *et al*., 2021) in line with attenuated mucosal expression of inflammatory cytokines (Shi *et al*., 2016). Patient findings are supported by data from animal models of colitis which show elevated Ang II in the colonic mucosa following induction of experimental colitis (Katada *et al*., 2008), and a reduction in inflammation, diarrhoea and mucosal pro-inflammatory cytokines following treatment with ACE inhibitors or ARBs (Spencer *et al*., 2007; Santiago *et al*., 2008; Shi *et al*., 2016). In addition, stimulation of Ang II production by renin overexpression, or chronic Ang II administration, precipitates colitis in mice and increased mucosal expression of TNFα, IL-1β, IL-6 and IL-17 (Shi *et al*., 2016).

Despite the emerging role for Ang II in colitis, its role in visceral nociception during colitis has not been extensively studied. To address this, we first demonstrated that Ang II stimulates nociceptors using *in vitro* Ca^2+^ imaging. Ang II elevated cytosolic Ca^2+^ in small diameter DRG neurons co-sensitive to the algogenic TRPV1 agonist capsaicin. Ang II-sensitive neurons also expressed Na_V_1.8, demonstrated by the co-localisation of Ca^2+^ transients with Na_V_1.8-tdTomato and the loss of Ang II-evoked Ca^2+^ transients following the ablation of Na_V_1.8-positive neurons. The direct activation of DRG neurons by Ang II was confirmed by repeating experiments following MACS to remove of non-neuronal cells (Thakur *et al*., 2014; Tewari *et al*., 2020) in which a comparable neuronal response to Ang II was observed. Given the loss of large-diameter neurons after MACS, one may expect an increase in the proportion of Ang II-sensitive neurons; this was not observed and may indicate a minor role for non-neuronal cells in the neuronal response to Ang II. Furthermore, this response was abolished by pre-treatment with valsartan, demonstrating that AT_1_ mediates the direct interaction between Ang II and nociceptors. Of note, the present study did not recapitulate the findings of Shepherd *et al*. (2018), who observed no increase in cytosolic Ca^2+^ in response to Ang II application to DRG sensory neurons (Shepherd *et al*., 2018). The reasons underpinning this discrepancy are not immediately clear, though differences in neuronal culture conditions are apparent, most notably a possible reduction AT_1_ receptor expression (Wangzhou *et al*., 2020) over the longer culture time used by Shepherd and colleagues. Nevertheless, our findings are consistent with functional studies in other cell types expressing AT_1_, including heterologous cell lines (Zitt *et al*., 1997), adrenal chromaffin cells (Liu *et al*., 2017) and central neurons (Gebke *et al*., 1998), which demonstrate AT_1_-mediated Ca^2+^ transients in response to Ang II.

TRPV1^+^ and Na_V_1.8^+^ neurons are known to mediate colonic hypersensitivity (Daou *et al*., 2016; Castro *et al*., 2019), indicating that their stimulation by AngII may contribute to hypersensitivity and pain in the inflamed bowel. Consistent with this, we demonstrated that Ang II elicited a marked increase in colonic afferent activity in tissue from both male and female mice. In keeping with the expression of AT_1_, but not AT_2,_ receptors in colonic neurons (Hockley *et al*., 2019), the Ang II mediated increase in colonic afferent activity was abolished by pre-treatment with AT_1_, but not AT_2_, antagonists. The response to Ang II could not be recapitulated by administration of an AT_2_ agonist. Furthermore, teased fibre recordings confirmed that the majority of colonic afferent fibres sensitive to Ang II were nociceptors based on their co-sensitivity to luminal distension at thresholds >20 mmHg. While AT_1_ is known to be expressed on colon-innervating neurons it is also expressed by other neurons (enteric neurons) and non-neuronal cells, such as enteroendocrine cells (Lu *et al*., 2019), within the gut. As such it is highly likely that the contribution of Ang II to visceral nociception in IBD may be mediated through a variety of neuronal and non-neuronal pathways alongside the direct activation of sensory neurons revealed in the present study. However, based on our data showing the loss of colonic afferent sensitivity to Ang II in tissue from mice in which the NP nociceptor population had been ablated, it would appear that the direct activation of AT_1_ expressing sensory neurons is a dominant pathway by which Ang II stimulates visceral nociceptors.

In summary, we have used genetic profiling of inflamed IBD tissue and sensory neurons to identify a novel pathway for visceral nociceptor activation in UC patients, namely Ang II/AT_1_-mediated activation of Na_V_1.8-positive colonic nociceptors. The findings from this study highlight the hitherto unrecognised potential to repurpose ARBs for the treatment of pain in UC.

## Supplemental information

### Supplementary methods and materials

#### Colonic biopsies

Colonic biopsies were taken following legal guardian consent from paediatric patients undergoing colonoscopy as part of their routine medical care. For patient data, see Supplemental Data 1. Biopsies were taken from patients undergoing diagnostic colonoscopy at the Royal London Hospital. Colonic biopsies were taken from sites of inflammation in UC patients, and CD patients divided into drug naïve (CDN) and treatment refractory (CDT) subgroups, with all groups reporting abdominal pain in the 4 weeks prior to endoscopy. Biopsies were also taken from the sigmoid colon of patients who reported symptoms of abdominal pain in the 4 weeks prior to endoscopy but showed no signs of inflammation on investigation and were subsequently diagnosed as having recurrent abdominal pain (RAP). Finally, biopsies were also taken from the sigmoid colon of non-inflamed control patients who reported no abdominal pain in the 4 weeks prior to colonoscopy and showed no evidence of inflammation following endoscopy. Ethical approval for the study was provided by the East London and The City Health Authority Research Ethics Committee (REC# P/01/023). Biopsy samples were collected in modified Krebs/HEPES buffer from which supernatants were taken for study in a separate series of experiments, following which samples were transferred to RNAlater and stored at -80°C until processing for RNA sequencing.

#### RNA sequencing of colonic biopsies

RNA was isolated from 30 mg of human colonic tissue using a RNeasy mini tissue kit (Qiagen), with DNase treatment. The resulting concentration of RNA was determined by NanoDrop 1000 (Thermo). RNA integrity was assayed with the Bioanalyzer (Agilent). Only RNA of suitable quality (i.e., RNA integrity number 8, rRNA ratio (28s/18s) 2, was used for RNA sequencing. Libraries were generated from 100 ng total RNA with NEBNext Ultra with polyA selection (NEB). RNA sequencing was performed at the Queen Mary University of London Genome Centre (https://www.qmul.ac.uk/blizard/genome-centre/) with Illumina NextSeq500 with an average of 44 million 75 bp paired end reads generated per sample. The quality of the sequencing reads (fastq files) was assessed by FastQC (version 0.11.2). The reads were trimmed for adaptor sequences and poor-quality reads with Trim Galore (version 0.3.7). Two trimming phases were applied, the first to remove adaptors and the second to remove poly G sequences. The quality of the trimmed sequences was reassessed with FastQC (version 0.11.2). After satisfactory quality control, trimmed sequences were aligned to the coding regions of the human reference genome (GRCh37) with TopHat2 (version 2.0.13) and bowtie2 (version 2.2.3). Transcript abundance was then calculated by HTSeq-counts software (version 0.6.0). Unadjusted transcript abundance was then exported to the R environment (version 3.1.2) for exploratory data analysis and differential expression analyses. The principal component analysis (PCA) and distance between samples from DESeq2 (version 1.6.3) were used to assess the dispersion and categorization of samples. Differential expression analysis was investigated with edgeR (version_3.8.6). Genes with low counts and expressed in only one sample per category were removed from further analysis. The calcNormFactors function was used to calculate the normalization factors to account for library sizes. Samples were then investigated for differences either between SR and AF or differences between the left and right atrium. Dispersion was calculated by using the functions estimateCommonDisp and estimateTagwiseDisp. The exact test was applied (exactTest) to obtain genes differentially expressed between non-inflamed, CD and UC biopsies.

#### Enrichment analysis

Enrichment analysis was carried out to identify signals of enriched cell types, signalling pathways, biological processes and subcellular compartments within the sets of genes upregulated in UC and CD (cut off, p < 0.0001) using EnrichR (https://maayanlab.cloud/Enrichr/). In Figures 1 and 2, the odds ratio is the abundance of genes corresponding to a particular enrichment term in the upregulated gene set compared to background. Significance of enrichment of particular terms was determined using Fisher’s exact test with Benjamini-Hochberg post-tests (terms were denoted as significantly enriched if p < 0.05). The combined score for an enrichment term is the product of the natural logarithm of the Benjamini-Hochberg-adjusted p-value and the z-score for the deviation from the expected rank of the term, providing further stratification of enriched terms.

#### Animals

All animal work was carried out in accordance with the Animals (Scientific Procedures) Act 1986 with prior approval under Home Office License PPL 70/7382. Mice were housed in cages of up to six littermates under a 12-hour light/dark cycle with enrichment (e.g., igloos and tunnels) and *ad libitum* access to food and water. Unless stated otherwise (see Figure 5J), all mice used were male aged 8-14 weeks on a C57Bl/6 background. Na_V_1.8^Cre^ (Jackson Laboratories stock 036564) (Nassar *et al*., 2004), ROSA26^CAG-flox-stop-tdTom^ (Jackson Laboratories stock 007905) (Madisen *et al*., 2010) and ROSA26^flox-stop-eGFP-DTA^ (Jackson Laboratories stock 032087) (Ivanova *et al*., 2005; Abrahamsen *et al*., 2008) mouse lines and genotyping protocols have been described previously. Tmem45b^Cre^ and Advillin^flox-stop-tdTom-DTA^ mouse lines were generated and characterised recently (Zhao *et al*., 2021).

#### Culture of sensory neurons from dorsal root ganglia

Dorsal root ganglia (DRG, T12-L6) were harvested and incubated with Lebovitz L-15 Glutamax media (Invitrogen) containing 1 mg/mL type 1A collagenase (Sigma Aldrich) and 6 mg/mL bovine serum albumin (BSA, Sigma Aldrich) for 15 min (37°C, 5% CO_2_). DRG were subsequently incubated with L-15 media containing 1 mg/mL trypsin (Sigma Aldrich) and 6 mg/mL BSA for 30 min (37°C, 5% CO_2_). DRG were gently triturated using a P1000 pipette tip and pelleted by centrifugation at 100*g* for 30 s. The supernatant (containing dissociated cells) was collected, and trituration was repeated five times. The collected supernatant was centrifuged at 100*g* for 5 min and pelleted cells were resuspended in L-15 media supplemented with 10% (v/v) foetal bovine serum, 2.6% (v/v) NaHCO_3_, 1.5% (v/v) D-glucose and penicillin/streptomycin and plated on to laminin- and poly-D-lysine-coated coverslips (MatTek). Cells were incubated at 37°C in 5% CO_2_ and were used for imaging after no more than 24 hours.

#### Ca^2+^ imaging of cultured sensory neurons

Cells were loaded with 10 µM Fluo-4-AM diluted in bath solution (in mM: 140 NaCl, 4 KCl, 1 MgCl_2_, 2 CaCl_2_, 4 D-glucose and 10 HEPES; pH 7.35-7.45) by incubation for 30-45 min (room temperature, shielded from light). Following incubation, coverslips were washed with bath solution and mounted on the stage of an inverted microscope (Nikon Eclipse TE2000S). For studies using antagonists, cells were pre-incubated with drug-containing solution (200 µL) for 10 min prior to imaging. During imaging, cells were superfused with bath solution at ∼0.5 mL/min using a gravity-fed perfusion tip.

Images were captured using a CCD camera (Retiga Electro, Photometrics) at 2.5 Hz with 100 ms exposure. Fluo-4 was excited by a 470 nm light source (Cairn Research) and emission at 520 nm was recorded using µManager. Where multiple drugs were added to the same coverslip, at least 3 min elapsed between applications. At the end of each experiment, 50 mM KCl was applied to identify viable neurons and enable the normalisation of fluorescence.

Image analysis was carried out using ImageJ. Regions of interest were manually drawn around cells and average pixel intensity per neuron was measured and analysed using custom-written scripts in RStudio. After the subtraction of background fluorescence, values were normalised to baseline fluorescence (10 s prior to drug application) and the maximal fluorescence during KCl application (F_pos_), such that 0 F/F_pos_ and 1 F/F_pos_ represent baseline and maximal fluorescence in KCl, respectively. Only cells which exhibited a rise in fluorescence of >5% over baseline during KCl application were included in analysis; no difference in the magnitude of the response to KCl was observed between experimental groups. Neurons were classed as responsive to a particular drug if fluorescence >0.1 F/F_pos_ was attained.

#### Magnetic-activated cell sorting (MACS) of cultured sensory neurons

DRG from 2−3 mice were isolated and cultured as above, but trypsin incubation was omitted and DRG were incubated with collagenase (1 mg/mL with 6 mg/ml BSA) for 45 min. Pelleted neurons were washed in 2 ml Dulbecco’s phosphate-buffered saline (DPBS, containing 0.9 mM CaCl_2_ and 0.5 mM MgCl_2_) and centrifuged for 7 min (100*g*). Pelleted cells were resuspended in MACS rinsing solution (120 μL, Miltenyi Biotec), supplemented with 0.5% w/v BSA (sterile filtered at 0.2 μM), and incubated (5 min at 4°C) with a biotin-conjugated non-neuronal antibody cocktail (30 μL, Miltenyi Biotec). DPBS was added to a volume of 2 ml and the suspension centrifuged for 7 min at 100*g*. The pellet was resuspended in 120 μL MACS rinsing solution with 30 μL biotin-binding magnetic beads (Miltenyi Biotec) and incubated for a further 10 min at 4°C, before being topped up to 500 μL with MACS rinsing buffer.

The cell suspension was filtered by gravity through a magnetic column (LD column, Miltenyi Biotec) primed with 2.5 mL MACS rinsing solution. Following the addition of the cell suspension, 1 ml MACS rinsing solution was used to collect the remnants of the cell suspension and passed through the column prior to a final wash. The 5 mL eluted was centrifuged for 7 min at 100*g* and the final pellet resuspended in supplemented L-15 medium, before plating on 35 mm poly-D-lysine-coated glass bottom culture dishes further coated with Matrigel (diluted 1:10 in L-15 medium). Cells were incubated in supplemented L-15 media at 37°C in 5% CO_2_ and were used for imaging after 48 hours.

#### Immunocytochemistry of cultured sensory neurons

DRG neurons were cultured as above and seeded onto 12 mm coverslips coated in poly-L-lysine and laminin. After 24−48 h in culture, cells were fixed at room temperature in 4% paraformaldehyde (10 min) and washed in PBS. Cells were permeabilized with 0.05% Triton-X100 for 5 min at room temperature. Cells were washed again in PBS and then blocking buffer (1% goat serum in 0.2% Triton-X100) was applied for 30 min. Cells were incubated with a rabbit anti-βIII-tubulin primary antibody (1:1000, Abcam: ab18207; RRID: AB_444319) for 3 hours at room temperature.

Following primary antibody incubation, cells were washed in PBS and incubated with an Alexa Fluor-568 goat anti-rabbit secondary antibody diluted in PBS (1:1000, Invitrogen: A11008; RRID: AB_143165) plus 4’-6-diamidino-2-phenylindole (DAPI; 1:1000, Abcam) for 1 hour at room temperature. After a final wash, coverslips were mounted, cell side down, on 25 × 75 × 1 mm glass slides using Mowoil 4−88 mounting medium (Sigma-Aldrich: 81381). Mounting medium was set at 4°C and slides were imaged within 2 hours.

Slides were imaged using an Olympus BX51 microscope. Fluorophores were excited with 568 nm (Alexa Fluor-568) or 350 nm (DAPI) light sources. Images were captured on a Qicam CCD camera (QImaging) with a 100 ms exposure and false coloured (βIII-tubulin, green; DAPI, blue). No βIII-tubulin staining was observed when the primary antibody was omitted (data not shown).

Images were analysed using ImageJ as previously described (Hartig, 2013). An automatic ‘minimum error’ threshold algorithm was applied to 8-bit images of βIII-tubulin or DAPI staining to distinguish background from objects of interest. Binary and raw images were manually compared, and the threshold manually adjusted to ensure all regions of interest were captured. The threshold was invariably placed within the first minimum after the major peak of the image histogram. Binary images then underwent watershed segmentation to separate distinct objects in close apposition. Identified objects, positive for βIII-tubulin and/or DAPI, were automatically counted using ImageJ and a ratio of βIII-tubulin-positive cells (neurons) to DAPI-positive cells (neurons and non-neuronal satellite cells) calculated.

#### Electrophysiological recording from the lumbar splanchnic nerve

For both multi- and single-unit recording, the colorectum (from splenic flexure to anus) with the associated lumbar splanchnic nerve was isolated and removed. The colorectum was flushed and transferred to a tissue bath before being cannulated and both luminally perfused (200 µL/min) and serosally superfused (7 mL/min; 32-34°C) with Krebs buffer (in mM: 124 NaCl, 4.8 KCl, 1.3 NaH_2_PO_4_, 25 NaHCO_3_, 1.2 MgSO_4_, 11.1 D-glucose and 2.5 CaCl_2_) supplemented with atropine (10 µM) and nifedipine (10 µM) to block smooth muscle activity. Luminal pressure was maintained between 2-5 mmHg (Neurolog NL108, Digitimer Ltd, UK). Activity from isolated bundles (or teased fibres) of the lumbar splanchnic nerve (rostral to the inferior mesenteric ganglia) was recorded using borosilicate glass suction electrodes. Signals were amplified (gain, 5 kHz), band-pass filtered (100-1500 Hz, Neurolog, Digitimer Ltd, UK) and digitally filtered for 50 Hz noise (Humbug, Quest Scientific, Canada). Signals were digitised (20 kHz, Micro1401, Cambridge Electronic Design, UK) and recorded using Spike2 (Cambridge Electronic Design, UK). Nerve discharge was quantified by determining the number of field potentials which were greater in magnitude than twice the background noise (typically 60-80 µV). Changes in nerve discharge were calculated by subtracting baseline firing (average of five minutes prior to drug application) from activity during drug application. In teased fibre experiments, single units were identified by waveform matching, allowing the properties of individual fibres to be determined (Hockley *et al*., 2016).

#### Statistics

All data were scrutinised to verify that they met the assumptions of parametric analyses. Normality was assessed using the Shapiro-Wilk test, and homogeneity of variances with F-tests; heterogeneity of variances was corrected using Welch’s correction where appropriate. Where the assumptions required for parametric analyses were not met, rank-based, non-parametric alternatives were used. Sample sizes were not prespecified before data acquisition, but inter-group comparisons were decided before data was obtained, and all statistical tests carried out are reported. Data are presented as mean ± standard error (SEM). P-value cut offs are denoted in figures as: *p<0.05, **p<0.01, ***p<0.001.

## Supplemental figures

**Supplemental Figure 1:**
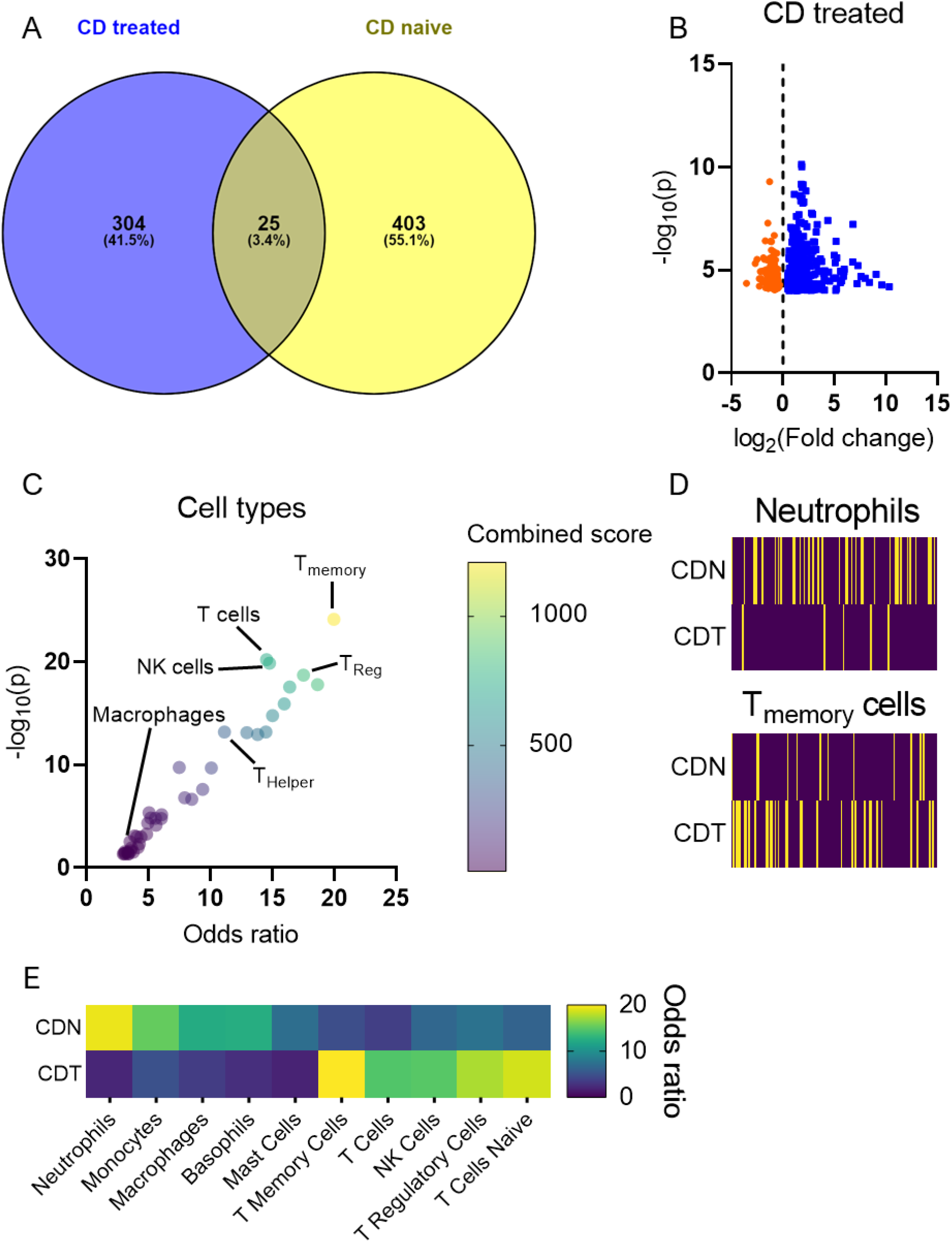
Differential gene enrichment in biopsies from CDN and CDT patients. (A) Venn diagram showing the proportion of shared and distinct genes elevated in CDN and CDT (relative to non-inflamed controls). (B) Volcano plot showing genes up- (blue) or down- (orange) regulated (p ≤ 0.0001) in CDT biopsies. (C) Bubble plots depicting the enrichment of gene ontology terms corresponding to different cell types in CDT biopsies. (D) Heatmaps showing the annotated gene sets used to identify neutrophils (top) and Tmemory cells (bottom); yellow indicates enrichment in biopsy tissue. Genes used to identify neutrophils are far less enriched in CDT tissue compared to CDN, while those used to identify Tmemory cells are more enriched in CDT. (E) Heatmap showing the odds ratio for the enrichment of gene ontology terms corresponding to particular immune cell types in biopsies from CDN and CDT patients.

**Supplemental Figure 2:**
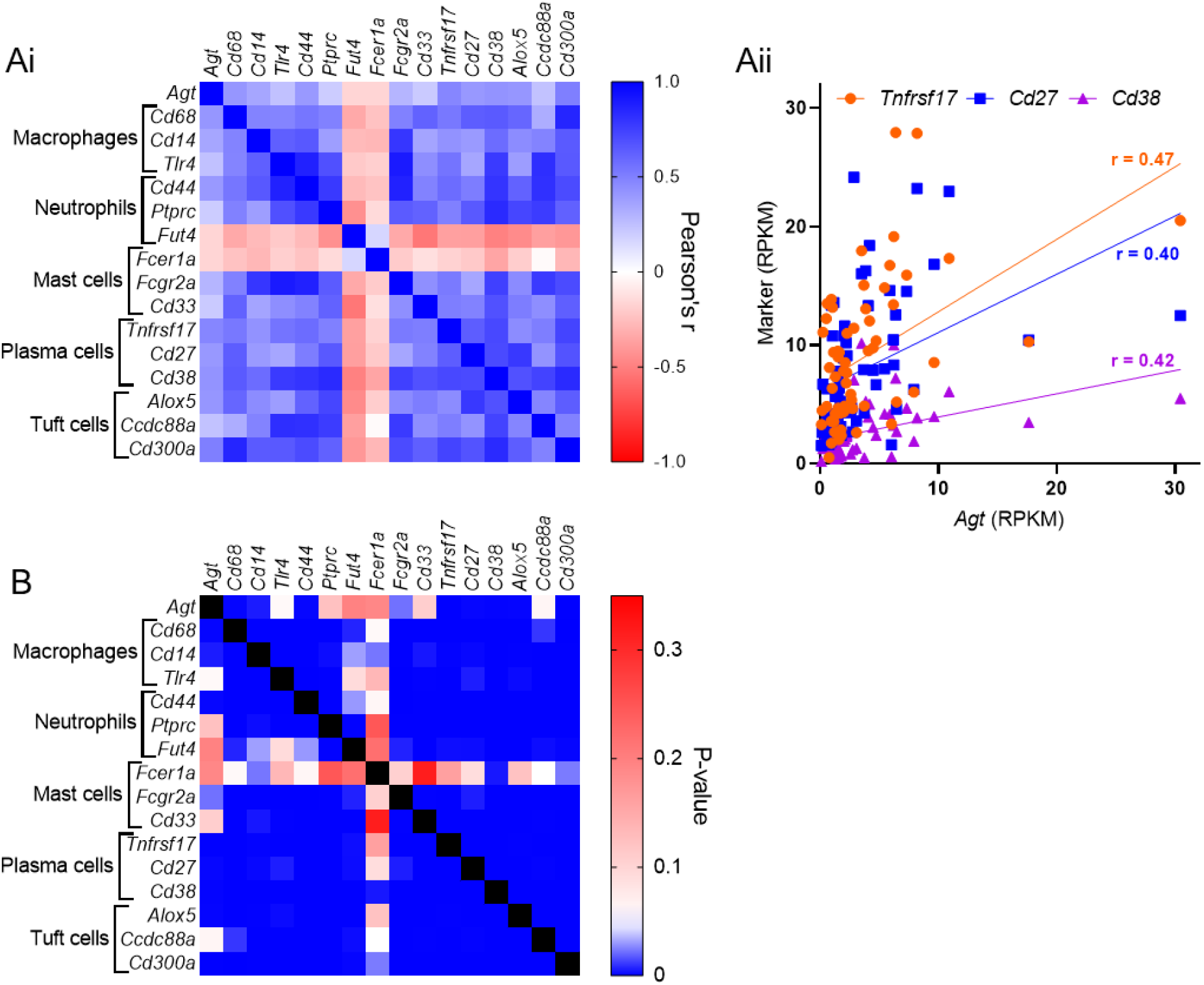
Potential sources of Agt. (A) (i) Correlation matrix for Agt and markers of cell types highly enriched in CD (macrophages and neutrophils) and UC (mast cells, plasma cells and tuft cells). Positive and negative correlations are shown with shades of blue and red, respectively. (ii) Example correlations between Agt and markers of plasma cells. All three markers exhibit a moderate positive correlation with Agt (Tnfrsf17: r = 0.47, p = 8.6×10-5; Cd27: r = 0.40, p = 0.001; Cd38: r = 0.42, p = 0.001). Data from patients in all groups included in analysis. (B) Matrix showing p-values for the correlations in (Ai). Red: p > 0.05, white: p = 0.05, blue: p < 0.05.

## Supplemental files

**Supplemental data 1:** Patient metadata.

**Supplemental data 2:** RNAseq analysis; sets of up- and down-regulated genes in CDN, CDT, UC and RAP/IBS patient groups.

**Supplemental data 3:** Enrichment analysis.

## Funding

This work was supported by the Biotechnology and Biological Science Research Council, LifeArc, Crohn’s and Colitis UK, The Gates Scholarship, The Cambridge Trust, Bowel Research UK and the British Society of Paediatric Gastroenterology, Hepatology and Nutrition.

## Disclosures

Dr Paul Wright is an employee of LifeArc.

No authors declare any conflict of interest, financial or otherwise.

## References

Abrahamsen, B. et al. (2008) ‘The cell and molecular basis of mechanical, cold, and inflammatory pain’, Science, 321(5889). doi: 10.1126/science.1156916.

Actis, G. C. et al. (2011) ‘Commonalities and differences between Crohn’s disease and ulcerative colitis: The genetic clues to their interpretation’, Inflammation and Allergy - Drug Targets. doi: 10.2174/187152811798104926.

Bautzova, T. et al. (2018) ‘5-oxoETE triggers nociception in constipation-predominant irritable bowel syndrome through MAS-related G protein–coupled receptor D’, Science Signaling, 11(561). doi: 10.1126/scisignal.aal2171.

Castro, J. et al. (2019) ‘Activation of pruritogenic TGR5, MRGPRA3, and MRGPRC11 on colon-innervating afferents induces visceral hypersensitivity’, JCI Insight, 4(20). doi: 10.1172/jci.insight.131712.

Chakrabarti, S. et al. (2020) ‘Human osteoarthritic synovial fluid increases excitability of mouse dorsal root ganglion sensory neurons: An in-vitro translational model to study arthritic pain’, Rheumatology (United Kingdom), 59(3). doi: 10.1093/rheumatology/kez331.

Chen, E. Y. et al. (2013) ‘Enrichr: Interactive and collaborative HTML5 gene list enrichment analysis tool’, BMC Bioinformatics, 14. doi: 10.1186/1471-2105-14-128.

Dabek, M. et al. (2009) ‘Luminal cathepsin G and protease-activated§ receptor 4: A duet involved in alterations of the colonic epithelial barrier in ulcerative colitis’, American Journal of Pathology, 175(1). doi: 10.2353/ajpath.2009.080986.

Daou, I. et al. (2016) ‘Optogenetic silencing of Nav1.8-positive afferents alleviates inflammatory and neuropathic pain’, eNeuro, 3(1). doi: 10.1523/ENEURO.0140-15.2016.

Fairbrass, K. M. et al. (2021) ‘Effect of ACE inhibitors and angiotensin II receptor blockers on disease outcomes in inflammatory bowel disease’, Gut. doi: 10.1136/gutjnl-2020-321186.

Garg, M. et al. (2020) ‘Imbalance of the renin-angiotensin system may contribute to inflammation and fibrosis in IBD: A novel therapeutic target?’, Gut, 69(5). doi: 10.1136/gutjnl-2019-318512.

Gebke, E. et al. (1998) ‘Angiotensin II-induced calcium signalling in neurons and astrocytes of rat circumventricular organs’, Neuroscience, 85(2). doi: 10.1016/S0306-4522(97)00601-5.

Gomez, R. A. et al. (1993) ‘Leukocytes synthesize angiotensinogen’, Hypertension, 21(4). doi: 10.1161/01.HYP.21.4.470.

Hartig, S. M. (2013) ‘Basic image analysis and manipulation in imageJ’, Current Protocols in Molecular Biology, (SUPPL.102). doi: 10.1002/0471142727.mb1415s102.

Hockley, J. R. F. et al. (2016) ‘P2Y receptors sensitize mouse and human colonic nociceptors’, Journal of Neuroscience, 36(8). doi: 10.1523/JNEUROSCI.3369-15.2016.

Hockley, J. R. F. et al. (2019) ‘Single-cell RNAseq reveals seven classes of colonic sensory neuron’, Gut, 68(4). doi: 10.1136/gutjnl-2017-315631.

Hültner, L. et al. (2000) ‘In Activated Mast Cells, IL-1 Up-Regulates the Production of Several Th2-Related Cytokines Including IL-9’, The Journal of Immunology, 164(11). doi: 10.4049/jimmunol.164.11.5556.

Ivanova, A. et al. (2005) ‘In vivo genetic ablation by Cre-mediated expression of diphtheria toxin fragment A’, Genesis, 43(3). doi: 10.1002/gene.20162.

Jacobs, J. D. et al. (2019) ‘Impact of Angiotensin II Signaling Blockade on Clinical Outcomes in Patients with Inflammatory Bowel Disease’, Digestive Diseases and Sciences, 64(7). doi: 10.1007/s10620-019-5474-4.

Jaszewski, R. et al. (1990) ‘Increased colonic mucosal angiotensin I and II concentrations in Crohn’s colitis’, Gastroenterology, 98(6). doi: 10.1016/0016-5085(90)91088-N.

Katada, K. et al. (2008) ‘Dextran sulfate sodium-induced acute colonic inflammation in angiotensin II type 1a receptor deficient mice’, Inflammation Research, 57(2). doi: 10.1007/s00011-007-7098-y.

Kobayashi, T. et al. (2020) ‘Ulcerative colitis’, Nature Reviews Disease Primers, 6(1). doi: 10.1038/s41572-020-0205-x.

Kuleshov, M. V. et al. (2016) ‘Enrichr: a comprehensive gene set enrichment analysis web server 2016 update’, Nucleic Acids Research, 44(1). doi: 10.1093/nar/gkw377.

Liu, C. H. et al. (2017) ‘Arrestin-biased AT1R agonism induces acute catecholamine secretion through TRPC3 coupling’, Nature Communications, 8. doi: 10.1038/ncomms14335.

Lu, V. B. et al. (2019) ‘Adenosine triphosphate is co-secreted with glucagon-like peptide-1 to modulate intestinal enterocytes and afferent neurons’, Nature Communications, 10(1). doi: 10.1038/s41467-019-09045-9.

Luiz, A. P. et al. (2019) ‘Cold sensing by Na V 1.8-positive and Na V 1.8-negative sensory neurons’, Proceedings of the National Academy of Sciences of the United States of America, 116(9). doi: 10.1073/pnas.1814545116.

Maddur, M. S. and Bayry, J. (2015) ‘B cells drive Th2 responses by instructing human dendritic cell maturation’, OncoImmunology, 4(5). doi: 10.1080/2162402X.2015.1005508.

Madisen, L. et al. (2010) ‘A robust and high-throughput Cre reporting and characterization system for the whole mouse brain’, Nature Neuroscience, 13(1). doi: 10.1038/nn.2467.

Mantaka, A. et al. (2021) ‘Is there any role of renin-angiotensin system inhibitors in modulating inflammatory bowel disease outcome?’, European Journal of Gastroenterology and Hepatology. doi: 10.1097/MEG.0000000000001912.

Massimino, L. et al. (2021) ‘The Inflammatory Bowel Disease Transcriptome and Metatranscriptome Meta-Analysis (IBD TaMMA) framework’, Nature Computational Science, 1(8). doi: 10.1038/s43588-021-00114-y.

Masuda, A. et al. (2002) ‘Th2 Cytokine Production from Mast Cells Is Directly Induced by Lipopolysaccharide and Distinctly Regulated by c-Jun N-Terminal Kinase and p38 Pathways’, The Journal of Immunology, 169(7). doi: 10.4049/jimmunol.169.7.3801.

Nassar, M. A. et al. (2004) ‘Nociceptor-specific gene deletion reveals a major role for Nav 1.7 (PN1) in acute and inflammatory pain’, Proceedings of the National Academy of Sciences of the United States of America, 101(34). doi: 10.1073/pnas.0404915101.

Owen, C. A. and Campbell, E. J. (1998) ‘Angiotensin II generation at the cell surface of activated neutrophils: novel cathepsin G-mediated catalytic activity that is resistant to inhibition.’, Journal of immunology (Baltimore, Md.: 1950), 160(3).

Richter, F. et al. (2012) ‘Interleukin-17 sensitizes joint nociceptors to mechanical stimuli and contributes to arthritic pain through neuronal interleukin-17 receptors in rodents’, Arthritis and Rheumatism, 64(12). doi: 10.1002/art.37695.

Roda, G. et al. (2020) ‘Crohn’s disease’, Nature Reviews Disease Primers, 6(1). doi: 10.1038/s41572-020-0156-2.

Santiago, O. I. et al. (2008) ‘An angiotensin II receptor antagonist reduces inflammatory parameters in two models of colitis’, Regulatory Peptides, 146(1–3). doi: 10.1016/j.regpep.2007.10.004.

Segond von Banchet, G., et al. (2013) ‘Neuronal IL-17 receptor upregulates TRPV4 but not TRPV1 receptors in DRG neurons and mediates mechanical but not thermal hyperalgesia’, Molecular and Cellular Neuroscience, 52. doi: 10.1016/j.mcn.2012.11.006.

Shepherd, A. J. et al. (2018) ‘Angiotensin II triggers peripheral macrophage-to-sensory neuron redox crosstalk to elicit pain’, Journal of Neuroscience, 38(32). doi: 10.1523/JNEUROSCI.3542-17.2018.

Shi, Y. et al. (2016) ‘Activation of the Renin-Angiotensin System Promotes Colitis Development’, Scientific Reports, 6. doi: 10.1038/srep27552.

Spencer, A. U. et al. (2007) ‘Reduced severity of a mouse colitis model with angiotensin converting enzyme inhibition’, Digestive Diseases and Sciences, 52(4). doi: 10.1007/s10620-006-9124-2.

Tewari, D. et al. (2020) ‘Granulocyte-macrophage colony stimulating factor as an indirect mediator of nociceptor activation and pain’, Journal of Neuroscience, 40(10). doi: 10.1523/JNEUROSCI.2268-19.2020.

Thakur, M. et al. (2014) ‘Defining the nociceptor transcriptome’, Frontiers in Molecular Neuroscience, 7(November). doi: 10.3389/fnmol.2014.00087.

Usoskin, D. et al. (2015) ‘Unbiased classification of sensory neuron types by large-scale single-cell RNA sequencing’, Nature Neuroscience, 18(1). doi: 10.1038/nn.3881.

Wangzhou, A. et al. (2020) ‘Pharmacological target-focused transcriptomic analysis of native vs cultured human and mouse dorsal root ganglia’, Pain, 161(7). doi: 10.1097/j.pain.0000000000001866.

Xie, Z. et al. (2021) ‘Gene Set Knowledge Discovery with Enrichr’, Current Protocols, 1(3). doi: 10.1002/cpz1.90.

Zeisel, A. et al. (2018) ‘Molecular Architecture of the Mouse Nervous System’, Cell, 174(4). doi: 10.1016/j.cell.2018.06.021.

Zhao, J. et al. (2021) ‘Tools for analysis and conditional deletion of subsets of sensory neurons’, Wellcome Open Research, 6. doi: 10.12688/wellcomeopenres.17090.1.

Zitt, C. et al. (1997) ‘Expression of TRPC3 in Chinese hamster ovary cells results in calcium-activated cation currents not related to store depletion’, Journal of Cell Biology, 138(6). doi: 10.1083/jcb.138.6.1333.

