## Supplemental information for "Transcriptomic profiling reveals a pro-nociceptive role for Angiotensin II in inflammatory bowel disease"

*Animals*

All animal work was carried out in accordance with the Animals (Scientific Procedures) Act 1986 with prior approval under Home Office License PPL 70/7382. Mice were housed in cages of up to six littermates under a 12-hour light/dark cycle with enrichment (e.g., igloos and tunnels) and *ad libitum* access to food and water. Unless stated otherwise, all mice used were male aged 8-14 weeks on a C57Bl/6 background. $Na_V1.8^{Cre}$ (Jackson Laboratories stock 036564) (Nassar *et al.*, 2004), $ROSA26^{CAG\text{-}flox\text{-}stop\text{-}tdTom}$ (Jackson Laboratories stock 007905) (Madisen *et al.*, 2010) and $ROSA26^{flox\text{-}stop\text{-}eGFP\text{-}DTA}$ (Jackson Laboratories stock 032087) (Ivanova *et al.*, 2005; Abrahamsen *et al.*, 2008) mouse lines and genotyping protocols have been described previously. $Tmem45b^{Cre}$ and $Advillin^{flox\text{-}stop\text{-}tdTom\text{-}DTA}$ mouse lines were generated and characterised recently (Zhao *et al.*, 2021).

**Supplemental figures**

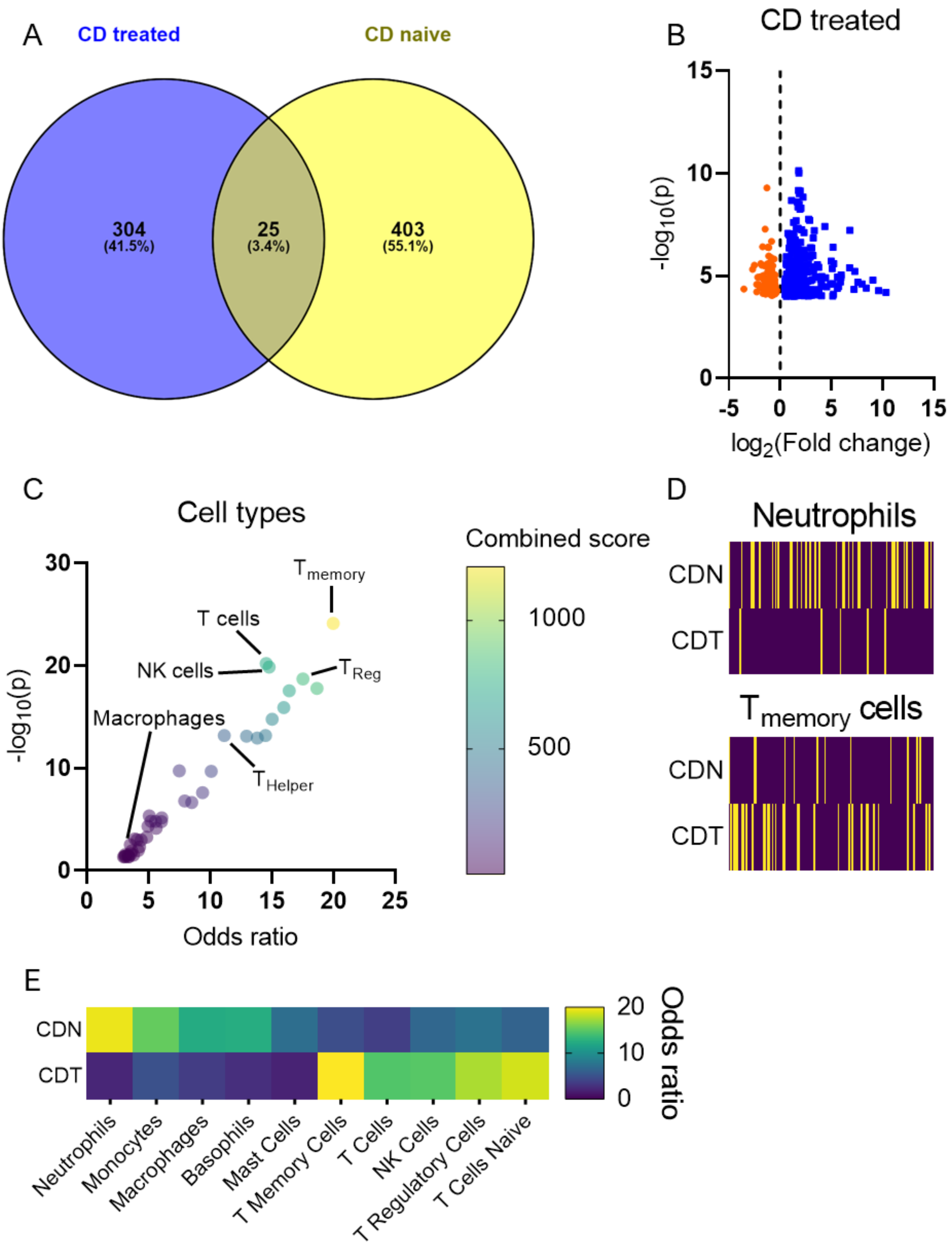


**Supplemental Figure 1: Differential gene enrichment in biopsies from CDN and CDT patients.**

(A) Venn diagram showing the proportion of shared and distinct genes elevated in CDN and CDT (relative to non-inflamed controls).

(B) Volcano plot showing genes up- (blue) or down- (orange) regulated ($p \leq 0.0001$) in CDT biopsies.

(C) Bubble plots depicting the enrichment of gene ontology terms corresponding to different cell types in CDT biopsies.

(D) Heatmaps showing the annotated gene sets used to identify neutrophils (*top*) and $T_{memory}$ cells (*bottom*); yellow indicates enrichment in biopsy tissue. Genes used to identify neutrophils are far less enriched in CDT tissue compared to CDN, while those used to identify $T_{memory}$ cells are more enriched in CDT.

(E) Heatmap showing the odds ratio for the enrichment of gene ontology terms corresponding to particular immune cell types in biopsies from CDN and CDT patients.

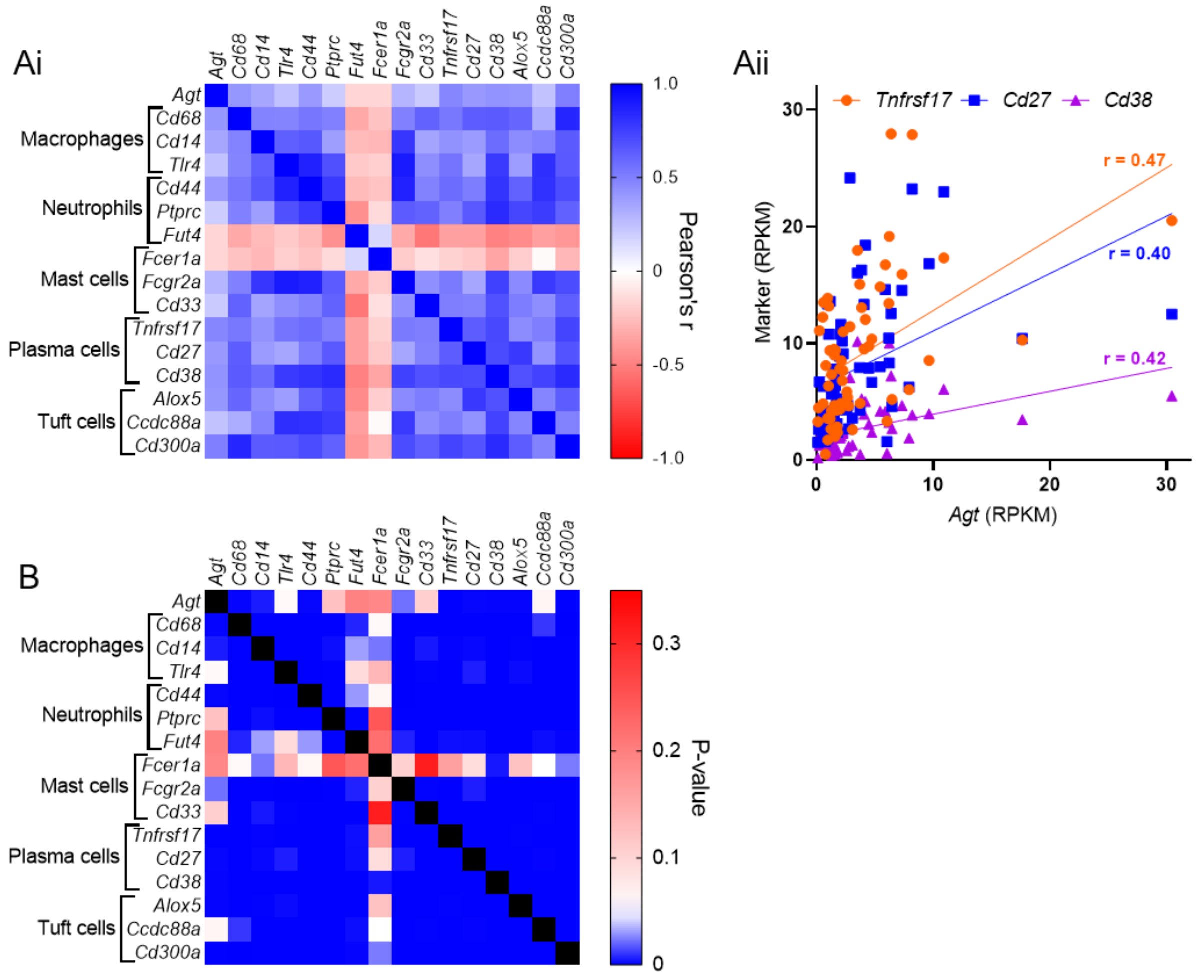


**Supplemental Figure 2: Potential sources of *Agt*.**

(A) *(i)* Correlation matrix for *Agt* and markers of cell types highly enriched in CD (macrophages and neutrophils) and UC (mast cells, plasma cells and tuft cells). Positive and negative correlations are shown with shades of blue and red, respectively. *(ii)* Example correlations between *Agt* and markers of plasma cells. All three markers exhibit a moderate positive correlation with *Agt* (*Tnfrsf17*: r = 0.47, p = $8.6 \times 10^{-5}$; *Cd27*: r = 0.40, p = 0.001; *Cd38*: r = 0.42, p = 0.001). Data from patients in all groups included in analysis.

(B) Matrix showing p-values for the correlations in (Ai). Red: $p > 0.05$, white: $p = 0.05$, blue: $p < 0.05$.

**Supplemental files**

**Supplemental data 1:** Patient metadata.

**Supplemental data 2:** RNAseq analysis; sets of up- and down-regulated genes in CDN, CDT, UC and RAP/IBS patient groups.

**Supplemental data 3:** Enrichment analysis.
