## Supplemental data 1 for "Transcriptomic profiling reveals a pro-nociceptive role for Angiotensin II in inflammatory bowel disease"

| | Age (SD), years | Sex F, % | N patients | N biopsies |
|---|---|---|---|---|
| **Non-inflamed** | 6.4 (3.4) | 37.5 | 8 | 14 |
| **CDN** | 12.6 (3.3) | 28.6 | 7 | 8 |
| **CDT** | 15.6 (1.4) | 42.9 | 7 | 11 |
| **UC** | 11.8 (4.9) | 44.4 | 9 | 9 |
| **Recurrent abdominal pain** | 12.0 (4.2) | 61.5 | 16 | 21 |

**Supplemental table 1: Metadata for each patient group.**

| | Sample | Prescribed treatment |
|---|---|---|
| **Non-inflamed** | 1 | NTR |
| | 2 | NTR |
| | 3 | Lactulose |
| | 4 | NTR |
| | 5 | NTR |
| | 6 | NTR |
| | 7 | NTR |
| | 8 | NTR |
| **CDN** | 1 | NTR |
| | 2 | NTR |
| | 3 | NTR |
| | 4 | NTR |
| | 5 | NTR |
| | 6 | NTR |
| | 7 | NTR |
| **CDT** | 1 | Azathioprine, mesalazine and infliximab |
| | 2 | Azathioprine and infliximab |
| | 3 | Azathioprine and mesalazine |
| | 4 | Azathioprine and mesalazine |
| | 5 | Azathioprine and prednisolone |
| | 6 | Azathioprine and infliximab |
| | 7 | Azathioprine and infliximab |
| **UC** | 1 | NTR |
| | 2 | NTR |
| | 3 | NTR |
| | 4 | NTR |
| | 5 | Azathioprine and mesalazine |
| | 6 | NTR |
| | 7 | Methotrexate and infliximab |
| | 8 | Metronidazole and ciprofloxacin |
| | 9 | NTR |
| **RAP** | 1 | NTR |
| | 2 | NTR |
| | 3 | NTR |
| | 4 | NTR |
| | 5 | NTR |
| | 6 | NTR |
| | 7 | NTR |
| | 8 | NTR |
| | 9 | NTR |
| | 10 | NTR |
| | 11 | NTR |
| | 12 | NTR |
| | 13 | Prednisolone |
| | 14 | NTR |
| | 15 | NTR |
| | 16 | NTR |

**Supplemental table 2: List of prescribed treatments for each patient.** NTR: no treatment.
